## Supplementary Information for "ORFanID: A Web-Based Search Engine for the Discovery and Identification of Orphan and Taxonomically Restricted Genes"

### SUPPLEMENTAL INFORMATION

**Richard S. Gunasekera[1*], Komal K. B. Raja[2], Suresh Hewapathirana[3], Emanuel Tundrea[4], Vinodh Gunasekera[5], Thushara Galbadage[6] and Paul A. Nelson[7]**

[1] Department of Chemistry, Physics and Engineering, School of Science, Technology & Health, Biola University, La Mirada, CA

[2] Department of Pathology & Immunology, Baylor College of Medicine, Houston, TX

[3] European Bioinformatics Institute, Welcome Genome Campus, Hinxton, Cambridgeshire, UK

[4] Griffiths School of Management and IT, Emanuel University of Oradea, Romania

[5] Bioinformatics, Chesalon USA, Inc., Houston TX

[6] Department of Kinesiology and Health Science, School of Science, Technology & Health, Biola University, La Mirada, CA

[7] Biola University, La Mirada, CA

*** Correspondence:**

Richard S. Gunasekera

**Keywords: ORFanID, orphan gene, ortholog, Taxonomically Restricted Genes (TRG), DNA, protein, bioinformatics**

**Supplemental Table 1.** Test results of ORFanID functionality.

| Species | Taxonomy ID | Accession | ORFanID Classification | Comments & References |
|---|---|---|---|---|
| Drosophila melanogaster | 7227 | NP_649722.1 | strict ORFan | Orphan gene based on flybase.org. Based on unpublished data, this protein has a role in eye development. |
| Drosophila melanogaster | 7227 | NP_727262.2 | strict ORFan | Orphan gene based on Levine et al., 2006. Flybase.org and ortho database show orthologs in Drosophila genus species. |
| Drosophila melanogaster | 7227 | NP_652406.1 | strict ORFan | Orphan gene based on Levine et al., 2006. No orthologs in orthodb. |
| Drosophila melanogaster | 7227 | NP_723266.1 | strict ORFan | Orphan gene based on Levine et al., 2006. No orthologs in orthodb. |
| Drosophila melanogaster | 7227 | NP_608405.4 | strict ORFan | *de novo* gene @ sub-group level based on Chen et al., 2007 & ortho db shows this protein in Drosophila sibling species, including *Drosophila melanogaster.* No orthologs in non-Drosophila insects or other organisms according to flybase.org |
| Drosophila melanogaster | 7227 | NP_726607.1 | Phylum | |
| Drosophila melanogaster | 7227 | NP_728356.2 | strict ORFan | Uncharacterized protein and no orthologs in flybase.org |
| Drosophila melanogaster | 7227 | NP_001097034.2 | strict ORFan | Reinhardt and Jones, 2013 |
| Drosophila melanogaster | 7227 | NP_001162900.1 | strict ORFan | Uncharacterized protein and no orthologs in flybase.org |

| | | | | |
|---|---|---|---|---|
| Drosophila melanogaster | 7227 | NP_001259080.1 | class | |
| C.elegans | 6239 | NP_500848.2 | order | TRG based on Verster et al., 2017. WormBase & orthodb show Nematode orthologs |
| C. elegans | 6239 | NP_494931.1 | orfan gene | TRG based on Verster et al., 2017. WormBase & orthodb show orthologs in Caenorhabditis genus |
| S.cervisiae | 4932 | NP_001335743.1 | strict ORFan | *de novo* gene. No orthologs in other databases |
| S. cervisiae | 4932 | NP_014130.1 | strict ORFan | Strict Orphan based on publication (Cai J et al., 2008).BSC4 may be involved in the DNA repair pathway during the stationary phase of S. cerevisiae and contribute to the robustness of S. cerevisiae, when shifted to a nutrient-poor environment. |
| S. cervisiae | 4932 | NP_015017.1 | strict ORFan | Essential TRG based on Verster et al., 2017 |
| S. cervisiae | 4932 | NP_010526.1 | strict ORFan | Essential TRG based on Verster et al., 2017 |
| S. cervisiae | 4932 | NP_011695.1 | strict ORFan | Essential TRG based on Verster et al., 2017 |
| S. cervisiae | 4932 | NP_011030.3 | strict ORFan | Essential TRG based on Verster et al., 2017 |
| S.cervisiae | 4932 | P47132.1 | strict ORFan | Strict Orphan based on publication. |
| S.cervisiae | 4932 | NP_009873.1 | strict ORFan | Saccharomyces specific gene based on few publications |

| | | | | |
|---|---|---|---|---|
| H. sapiens | 9606 | NP_073152.2 | Genus | Order restricted based on a publication and family based on ortho database |
| H. sapiens | 9606 | NP_001316897.1 | order | Order restricted based on a publication. Orthodb shows conservation in eutheria clade |
| H. sapiens | 9606 | NP_872390.2 | order | It is a human-specific gene based on a publication. No conserved domains in NCBI database. Eutheria clade based on orthodb |
| H. sapiens | 9606 | Q1W209.2 | family | It is a human-specific gene based on a publication |
| Danio rerio | 7955 | XP_002665008.4 | ORFan gene | It was shown to be genus restricted. According to orthodb, this protein has orthologs in actinopterygii class |
| Danio rerio | 7955 | NP_571379.1 | class | Should be present in all seeing animals. should be multi-domain |

**Supplemental Table 2.** abSENSE vs. ORFanID comparison.

| Organism | NCBI Protein Accession Number | ORFanID Classifier | Result |
|---|---|---|---|
| Fungal | NP_015174.1 | Class | 1676102403557_GlL |
| Fungal | NP_116703.5 | **Strict ORFan** | 1676139596145_gQr |
| Fungal | NP_012059.3 | **Strict ORFan** | 1676139596145_gQr |
| Fungal | NP_011401.1 | Genus | 1676139596145_gQr |
| Fungal | NP_014025.1 | Kingdom | 1676139596145_gQr |
| Fungal | NP_010070.1 | **Strict ORFan** | 1676139596145_gQr |
| Fungal | NP_013577.1 | Family | 1676139596145_gQr |
| Fungal | NP_010143.1 | Family | 1676139596145_gQr |
| Fungal | NP_010972.1 | Family | 1676139596145_gQr |
| Fungal | NP_010486.3 | Genus | 1676141964287_UiA |
| Fungal | NP_012181.3 | **Strict ORFan** | 1676141964287_UiA |
| Fungal | NP_014964.1 | Genus | 1676141964287_UiA |
| Fungal | NP_009719.3 | Family | 1676141964287_UiA |
| Fungal | NP_010398.3 | **Strict ORFan** | 1676141964287_UiA |
| Fungal | NP_014522.1 | Family | 1676141964287_UiA |
| Fungal | NP_012054.1 | **Strict ORFan** | 1676141964287_UiA |
| Fungal | NP_010175.1 | Phylum | 1676141964287_UiA |
| Fungal | NP_010486.3 | Genus | 1676143078659_3qE |
| Fungal | NP_009757.1 | Family | 1676143078659_3qE |
| Fungal | NP_012587.3 | **Strict ORFan** | 1676143078659_3qE |
| Fungal | NP_015256.1 | **Strict ORFan** | 1676143078659_3qE |
| Fungal | NP_013688.1 | Kingdom | 1676143078659_3qE |
| Fungal | NP_010017.2 | Genus | 1676143078659_3qE |
| Fungal | NP_013113.1 | Kingdom | 1676143078659_3qE |

| Organism | NCBI Protein Accession Number | ORFanID Classifier | Result |
|---|---|---|---|
| Fungal | NP_015111.1 | Genus | 1676143078659_3qE |
| Fungal | NP_116686.3 | Genus | 1676144099201_5oG |
| Fungal | NP_010885.1 | Genus | 1676144099201_5oG |
| Fungal | NP_014908.1 | Genus | 1676144099201_5oG |
| Fungal | NP_012282.3 | **Strict ORFan** | 1676144099201_5oG |
| Fungal | NP_013614.1 | Genus | 1676144099201_5oG |
| Fungal | NP_013929.1 | Family | 1676144099201_5oG |
| Insect | NP_001285241.1 | **Strict ORFan** | 1676207776266_kNn |
| Insect | NP_001097257.1 | **Strict ORFan** | 1676207776266_kNn |
| Insect | NP_731242.1 | **Strict ORFan** | 1676207776266_kNn |
| Insect | NP_572277.2 | Order | 1676207776266_kNn |
| Insect | NP_001163736.1 | **Strict ORFan** | 1676207776266_kNn |
| Insect | NP_001260070.1 | Family | 1676207776266_kNn |
| Insect | NP_001163602.1 | Phylum | 1676207776266_kNn |
| Insect | NP_001247103.1 | Phylum | 1676210851491_1sk |
| Insect | NP_648151.2 | Order | 1676210851491_1sk |
| Insect | NP_650536.1 | Family | 1676210851491_1sk |
| Insect | NP_001188817.2 | Order | 1676210851491_1sk |
| Insect | NP_611851.1 | Family | 1676210851491_1sk |
| Insect | NP_001262507.1 | Order | 1676210851491_1sk |
| Insect | NP_001137829.1 | Genus | 1676210851491_1sk |
| Insect | NP_996411.2 | Class | 1676232140982_SDo |
| Insect | NP_650020.2 | Family | 1676232140982_SDo |
| Insect | NP_572410.2 | **Strict ORFan** | 1676232140982_SDo |
| Insect | NP_612110.1 | **Strict ORFan** | 1676232140982_SDo |

| Organism | NCBI Protein Accession Number | ORFanID Classifier | Result |
|---|---|---|---|
| Insect | NP_001097941.1 | **Strict ORFan** | 1676232140982_SDo |
| Insect | NP_788714.1 | Family | 1676232140982_SDo |
| Insect | NP_523663.1 | Phylum | 1676232140982_SDo |
| Insect | NP_651860.2 | **Strict ORFan** | 1676232140982_SDo |
| Insect | NP_001246258.1 | Order | 1676234237302_RAr |
| Insect | NP_001262397.1 | Order | 1676234237302_RAr |
| Insect | NP_001286126.1 | Order | 1676234237302_RAr |
| Insect | NP_570044.1 | **Strict ORFan** | 1676234237302_RAr |
| Insect | NP_728059.1 | **Strict ORFan** | 1676234237302_RAr |
| Insect | NP_001287054.1 | Family | 1676234237302_RAr |
| Insect | NP_649652.4 | **Strict ORFan** | 1676234237302_RAr |
| Insect | NP_001285134.1 | Class | 1676234434226_fi4 |
